# Merkel cell polyomavirus regulates miR183 cluster and piR62011 in Merkel cell carcinoma

**DOI:** 10.1101/2022.01.18.476776

**Authors:** Reety Arora, Lamiya Dohadwala, Bageshri Naimish Nanavati, Vairavan Lakshmanan, Sanjukta Mukherjee

**Author notes:** shared first author (Reety Arora, Lamiya Dohadwala, Bageshri Naimish Nanavati).

## Abstract

Merkel cell carcinoma (MCC) is a rare, aggressive skin cancer, a major subset of which is caused by the clonal integration of Merkel Cell Polyomavirus (MCV). Recent studies by Cheng et al. (2017) reported that virus-derived small T antigen protein-bound EP400 complex drives expression of genes essential for cellular transformation. On close analysis of their ChIP-Seq data, we uncovered that the complex binds to the promoter region of the microRNA-183 cluster. The miRNA183 cluster is a cluster of 3 miRNAs (miR183, 182 & 96) expressed and regulated together. These miRNAs are conserved across species, highly expressed in human embryonic stem cells and necessary for sensory/ mechanosensory organ development. We hypothesized that the MCV oncoproteins regulate host miRNA expression directly; an interaction novel in polyomaviruses. We tested miRNA expression via qPCR in both virus positive and negative MCC cell lines and found the former showed a much higher level. Further, fibroblasts expressing T antigens displayed an increase in miR182 expression in comparison to control. Knock-down of T antigens in MCC cells correspondingly decreased miR182 levels. To investigate its regulation we performed luciferase assays for the miRNA predicted promoter that showed increased activity in the presence of T antigens. Intriguingly, the seed sequence of miR182 completely matches to a piRNA called piR62011. Upon reanalysis of a MCC small RNA library, piR62011 emerged as the highest expressed. We found it expressed in MKL-1, a MCV positive cell line as well. Finally, to translate our findings into therapy for MCC, we screened small molecule (CMBL) library by performing surface plasmon resonance (SPR) assay and identified small molecules that binds to pre-miRNA182 and are testing them for their activity to kill MCC.

## INTRODUCTION

Merkel cell carcinoma (MCC) is a rare and aggressive primary cutaneous malignancy with a poor prognosis and a mortality rate 33-46% greater than that of melanoma [1]. Sequencing studies of MCC samples revealed that there exists two different subtypes of MCC. One subtype was characterised by large number of somatic mutations typically caused by UVB radiation and the other showed fewer mutations with clonal integration of the Merkel cell polyomavirus (MCV) genome. Virus positive (MCV+) MCC reported in 80% of MCC cases show integration of MCV genome encoding viral transforming genes, large and small tumor antigens (LT and ST) required for tumor initiation and maintenance[1-3].

In all virus-positive MCC cases reported to date, LT has undergone truncations that disrupt viral replication activities but leave the LXCXE, RB-binding motif intact, thereby, allowing entry into the S phase of the cell cycle facilitating carcinogenesis[4]. The functional domains of ST harbours the protein phosphatase 2A (PP2A) binding site and the LT stabilising domain (LSD). LSD regulates the levels of LT by inhibiting (Fbw7) F-box and WD repeat domain-containing 7 of the ubiquitin ligase complex thus preventing the proteasomal degradation of LT [5, 6]. In addition to Fbw7, LSD is required for hyperphosphorylation of eukaryotic translation initiation factor 4E-binding protein 1 (4E-BP1) which represses its inhibitory action during translation regulation[7]. To further understand how MCV ST drives oncogenesis, Cheng *et. al*. (2017) demonstrated that ST-MYCL-EP400-MAX complex binds to the promoter region of several genes to activate gene expression in MCV positive MCC cell lines[4].

MicroRNAs (miRNA) are a group of small non-coding RNA which are 22-24 nucleotides in length and are involved in degradation of targets and translational repression of coding messenger RNAs by sequence-specific complementarity [8]. Since their discovery in the early 1900s, microRNAs have emerged as key players in post-transcriptional gene regulation [8]. MiRNAs are transcribed by RNA polymerase II from the genomic loci as nascent primary transcript (pri-miRNA) which is ∼110 nucleotides in length. Pri-miRNAs consisting of stem-loop structures are cleaved by an RNase III-like enzyme called Drosha in a microprocessor complex to yield a precursor transcript. The pre-miRNA is trafficked to the cytoplasm where it undergoes a Dicer mediated processing to give rise to an miRNA duplex consisting of a guide strand and passenger strand, where the latter is degraded and the mature guide strand is loaded onto the RNA-induced silencing complex (RISC), primarily, composed of Argonaute (Ago) protein. The guide strand binds to the 3’ untranslated region (UTR) of the target mRNA by complementary base pairing and is subjected to cleavage by Ago[9].

The genomic location of the miRNAs may be intergenic, intronic (in a special case of mirtons, the miRNA spans the intron completely and is flanked by splice sites on either ends) or exonic (rarely) [8, 9]. The miRNAs may be monocistronic i.e. transcribed from a single promoter or may occur in clusters, usually in less than 10 Kb, and are transcribed in a polycistronic fashion.

Despite all the information gathered in the past two decades on the tissue-specific expression of microRNAs and their aberrant expression patterns in a multitude of cancers, the mechanisms regulating their own expression remain largely unknown.

In this study, we sought to analyse the transcriptional regulation of miR183 cluster in the context of a rare and highly aggressive skin cancer, Merkel cell carcinoma. MicroRNA183 cluster is a large intergenic miRNA gene family comprising of 3 miRNA sequences namely miR183, miR96 and miR182 which are paralogous in nature. It is a highly conserved miRNA group mapped on the q arm of chromosome 7 (7q32.2) which transcribes in a polycistronic fashion[10]. The intergenic region between miR96 and miR182 spans a length of 4.3kb, however the spacing between miR183 and miR96 is shorter of 321bp (Sequence from NCBI, GRCh38.p12). The miR183 family exercises an important role in the development and maturation of sensory organs through a synchronised expression of the 3 miRNAs. This cluster is expressed in embryonic stem cells, and in the inner ear[11] and retina[12]. It was also observed that a temporal reduction in the expression of miR183 cluster is necessary for epidermal differentiation to neuroectodermal lineage[13].

Here, we describe that the small T antigen of Merkel cell polyomavirus is involved in the transcriptional regulation of miR183 cluster in Merkel cell carcinoma. Dual reporter luciferase assay confirmed the binding of MCV ST to the miR183 cluster promoter in MCV positive MCC cell line expressing T antigens versus in the absence of T antigens. In addition, we demonstrated the potential of novel Cyclic Mismatch Binding Ligands(CMBL)[14-16] molecules as future therapy for MCC due to its specificity and inhibitory action towards pre-miR182 maturation.

In sum, our study provides new insights into the regulation of this cluster in MCC development and opens up new opportunities for therapeutic intervention using cyclic mismatch binding ligands (CMBLs).

## RESULTS AND DISCUSSIONS

Ning et al. (2014)[17] characterised the miRNome of Merkel cell carcinoma (MCC) by next-generation sequencing (NGS) of small RNA libraries. They had identified that MCC shows an upregulation of miR183 cluster when compared to normal skin and other cutaneous tumors. We reanalyzed the small RNA libraries and plotted both 5p and 3p reads for all miRNA especially for the miRNA183 cluster – miRNA183, miRNA182 and miRNA96 (Fig 1 A and B). We found a differential fold expression between MCC_Rep1 and MCC_Rep2 (the two replicates studied in the original Ning et al. study)[17]. MS1, an MCV positive MCC cell line had similar expression to that seen in MCC_Rep1. We hypothesized that MCC_Rep1 was possibly virus positive, similar to MS1 and MCC_Rep2 was possibly virus negative. Unfortunately, we could not test this in the original samples, as they had been used entirely for the previous study.

**Fig 1.**
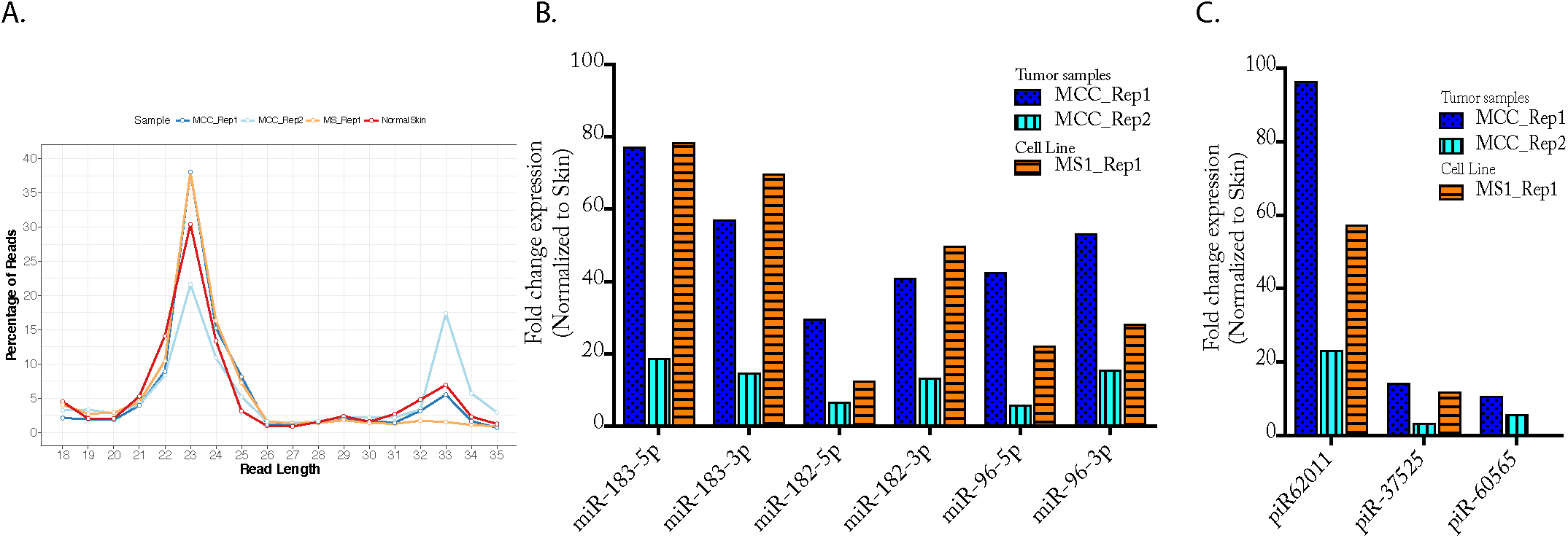
MiRNA183 cluster and piRNA62011 shows differential expression in Merkel tumors. **A**. Percent reads in MCC miRNome dataset from Ning et al., 2014 was plotted against read length. Two peaks, one at 21-24bp and another ar 32-34bp was observed. **B**. miRNA183 cluster (all 3 miRNAs, both 5p and 3p) was analyzed as fold change expression when normalized to skin. MCC_Rep1 and MS_1Rep1 showed similar values whereas MCC_Rep2 did not. **C**. piRNA expression was analyzed for each of the MCC datasets and when normalized to skin piRNA62011 showed a 96.5, 23.4 and 57.4 fold increase for MCC_Rep1, MCC_Rep2 and MS1_Rep1 respectively

Upon further evaluation of the small RNA library data, we found a peak at 32-34bp for the samples. We investigated this further and found that they all mapped to piRNAs (http://pirnabank.ibab.ac.in/) (Fig 1A). One piRNA, piR62011 stood out as it showed a fold increase of 96.5, 23.4 and 57.4 for MCC_Rep1, MCC_Rep2 and MS1_Rep1 respectively (Fig 1C). Although the other piRNAs did also show an MCC specific fold increase, it as not as substantial (14 fold compared to 96 fold, Fig 1C). piR62011 is a unique piRNA as its shares its sequence entirely with miRNA182[18]. Both piR62011 and miR182 have common seed sequences and we found them to be upregulated in MCC.

To further test our hypothesis, we established a qPCR method to detect miRNAs in total RNA from MCC cell lines and correlate Ct values to copy number via standard curves (Fig 2A). We found that all 3 miRNAs show much higher expression in MCV positive MCC cells as compared to MCV negative cells although the copy numbers vary per miRNA, per cell line (Fig 2B).

**Fig 2.**
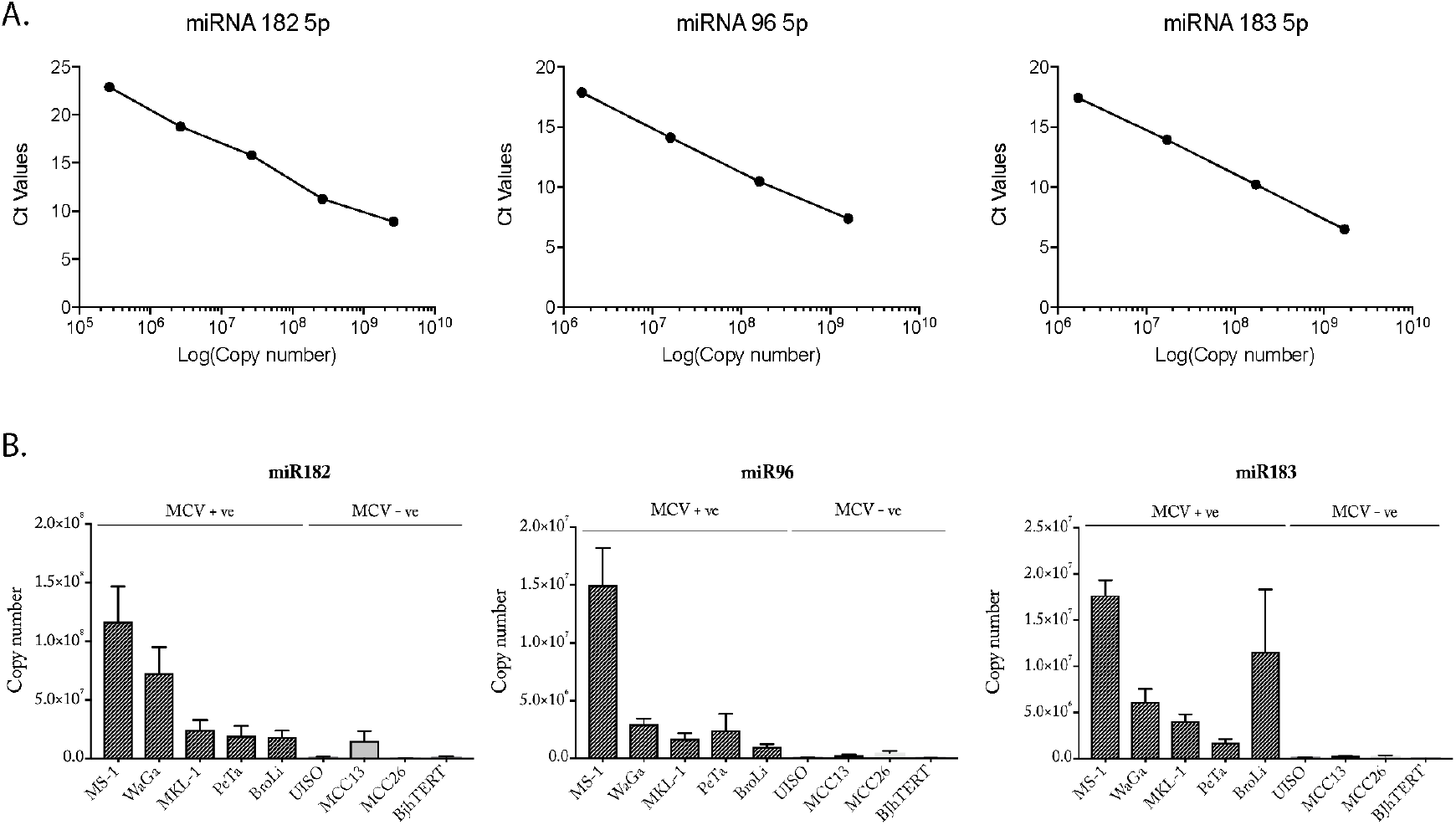
MiRNA183 cluster shows higher expression in MCV positive Merkel tumors. **A**. Standard curves for miRNA182, 96 and miR183 were made for qPCR Ct vs copy number. **B**. Plots showing Copy number of the 3 different miRNAs (miR182,miR96 and miR183) in 5 MCV positive MCC cell lines (MS-1, WaGa, MKL-1, PeTa and BroLi), 3 MCV negative MCC cell lines (UISO, MCC13 and MCC26) and BjhTERT fibroblasts. All 3 miRNAs show much higher expression in MCV positive MCC cells as compared to MCV negative cells although the copy numbers vary per miRNA, per cell line.

To understand why miRNA183 cluster is high in MCV positive MCC we analyzed ChIP seq data for MKL-1 cells chromatin-immunoprecipitated using H3K4me3, MAX, EP400 and ST from Cheng et al 2017 [4], using UCSC browser. A clear peak of H3K4me3 that also overlayed with MAX, EP400 and ST was seen 5222bp from the miR183 sequence on chromosome 7 on genomic DNA (Fig 3A). H3K4me3 is an activation mark often seen on promoter regions of genes[19]. A recent study defined the miRNA183 cluster promoter on chromosome 7, ∼5000bp upstream from miR183[20]. We tested 431bp of this region in a luciferase assay in the presence and absence of MCV T antigen expression. MCV T antigens show a 2.86 fold increase when compared to control (Fig 3B), further suggesting that T antigens transcriptionally up-regulate miRNA183 cluster its promoter.

**Fig 3.**
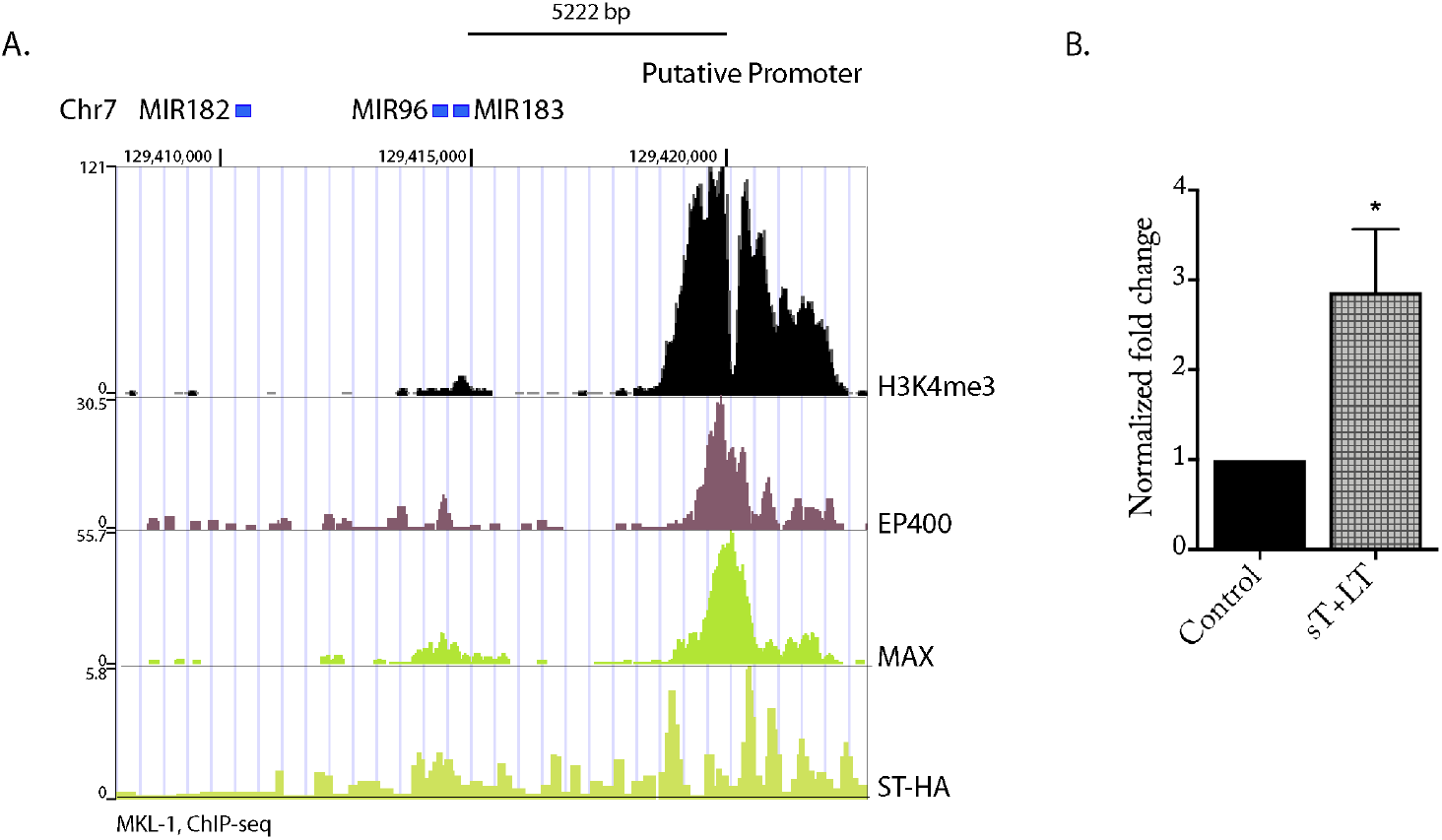
MiRNA183 cluster is regulated by MCV T antigens. **A**. ChiP –Seq Analysis of MKL-1 cells chipped using H3K4me3, EP400, MAX and ST-HA (overexpressed) from Cheng et al., 2017 shows a peak at a distance of 5222bp from miR183 sequence on chromosome7. This is the predicted, putative promoter for this cluster of miRNA. Peaks of H3K4me3 (activation mark) correspond with EP400, MAX and ST. **B**. Luciferase assay for the miRNA183 cluster putative promoter in the presence and absence of MCV T antigens shows a clear increase in Luciferase activity when T antigens were expressed. pGL4 vector expressing 431bp of the putative promoter was used for the experiment. Firefly luciferase was normalized to renilla luciferase (used as a transfection control) and then relative luciferase unit values were normalized to control.

Sequence-specific recognition of pre-miRNA by small molecules to repress the expression of miRNAs dysregulated in cancer are promising chemotherapeutic agents. Mukherjee *et al* (2016)[14] designed a library of compounds called cyclic mismatch binding ligands (CMBLs) containing two nucleobase recognition heterocyclic moieties linked by two variable linkers in cyclophane scaffold. Total 21 CMBL molecules[21] comprising two 2-amino-1,8-naphthyridine were developed[14-16] to target consecutive guanines (Gs) in DNA and RNA. Expression of 41 miRNAs expressing consecutive Gs in their pre-miRNA loop regions revealed miRNA182 to be highest and thus a valid target in MCC (Fig 4C). An *in vitro* Dicer-cleavage assay and Surface Plasmon Reasonance study of these 21 molecules helped us identify CMBL 3aL and 4a for further testing (Fig 5A, B and C). CMBL3aL and 4a showed notable response in SPR with rapid association kinetics, however CMBL3aL exhibited faster dissociation kinetics than CMBL4c.

**Fig 4.**
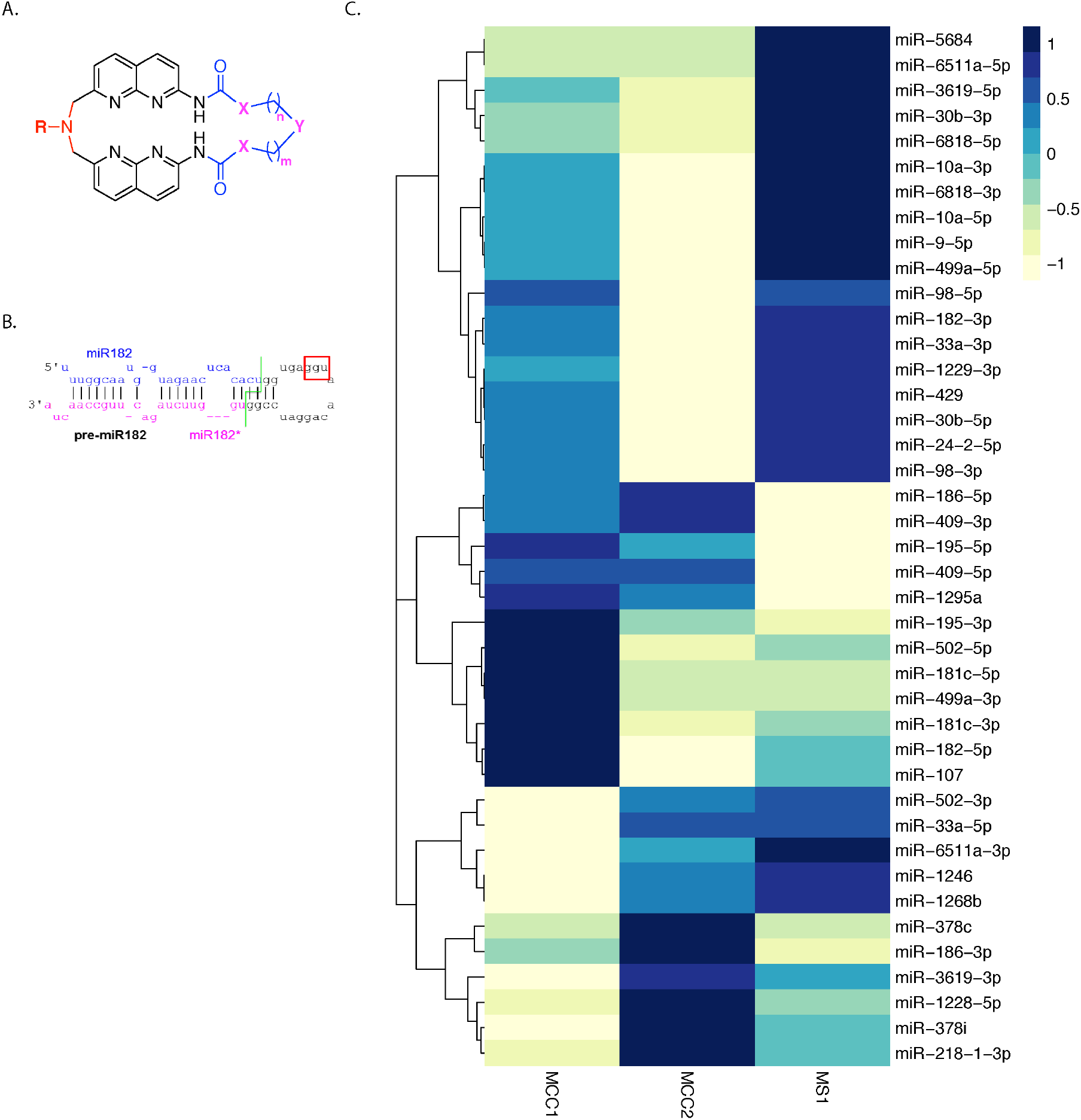
CMBLs can recognize consecutive guanine (GG) in hairpin loop of pre-miRNAs to inhibit its function. **A**. General structure of Cyclic Mismatch Binding Ligands (CMBL). R is H or or L [= (CH_2_)_3_NHCO(CH_2_)_3_NH_2_]; X = CH_2_ or O; Y = *E*-(CH=CH), *Z*-(CH=CH), CH_2_, NH and O; n, m = 1, 2 and/or 3. The linkers can be altered to create variations with target specificity. We are testing 21 such molecules that can target consecutive guanine (GG) sequences. **B**. Structure of pre-miR182 reveals presence of consecutive guanine (GG) sequence in loop and hence it can be a target for CMBL molecules. **C**. Heatmap depicts the expression of 41 miRNAs out of 1235 miRNAs in MCC that containing expected target motif for CMBLs in their precursor (pre-miRNA). Dataset from -small RNA library-Ning eta l 2013.

**Fig 5.**
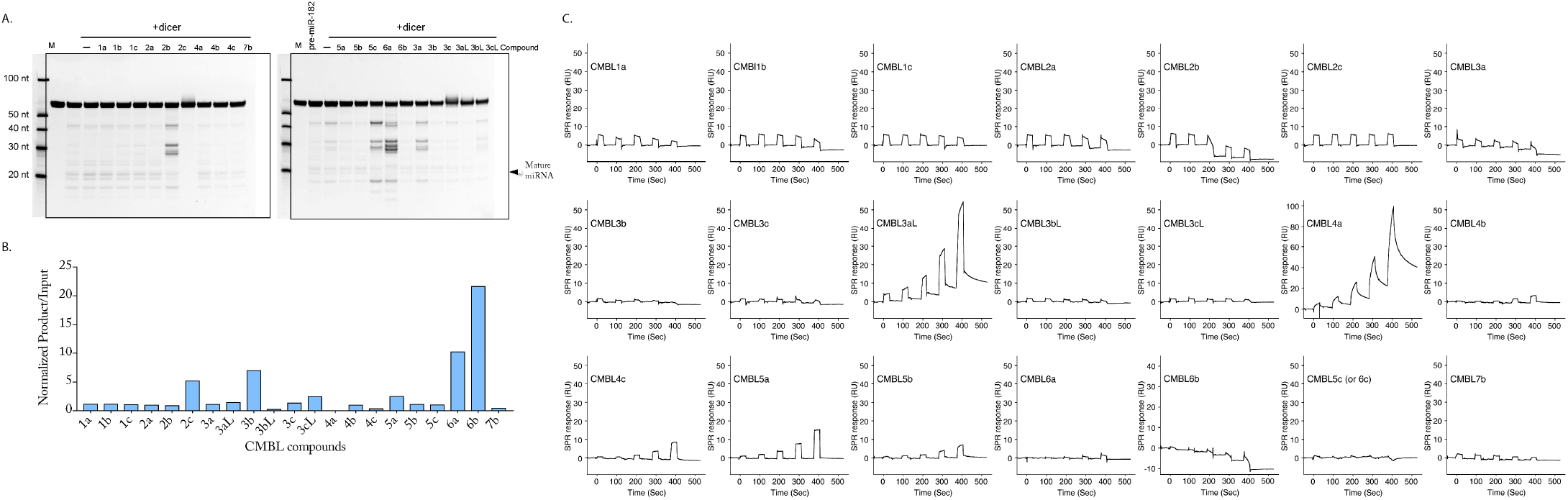
CMBLs can inhibit cleavage of pre-miRNA182. **A**. In vitro Dicer processing assay analysis. **B**. Densitometric analysis of in vitro dicer processing of pre-miR182 in the presence of CMBL molecules. This data revealed that out of the 21 analogues, CMBL 4a showed the least (0.03 unit) degree of Dicer cleavage indicating greater potency of cleavage inhibition. **C**. SPR analysis of 21 CMBLs with pre-miR182 (557.3 RU) immobilized on a sensor surface. The peaks in the graph correspond to the effect of 0.25, 0.5, 1.0, 2.0 ad 4.0 μM concentration of CMBLs.

Further testing of these CMBL molecules for their ability to inhibit miRNA182 expression in MCC and whether this reduction can be exploited for MCC therapy is ongoing. They serve as promising candidates for the Merkel tumor therapy and further testing will analyze them for the same.

## MATERIALS AND METHODS

### Bioinformatic analysis

We used Hg19(8) as the reference genomes (http://genome.ucsc.edu/) for this analysis. From the sequencing reads, we trimmed TruSeq small RNA adapters using customized perl script and cutadapt program(9). The adapter trimmed reads were aligned to rRNA and unaligned reads were taken for further analysis. We then segregated reads that are 18-35 nucleotides and mapped to genome and miRNA (http://www.mirbase.org/), tRNA (http://gtrnadb.ucsc.edu/), piRNA (http://pirnabank.ibab.ac.in/) databases using bowtie v1.0.0 (10). We used 2 mismatches as a constant parameter for all the alignments done in this study, as we did not observe much deviation in our results with varying mismatches ranging from 0-2. We used DESeq (11) for normalization and identification of differentially expressed small RNAs (adj. P-value < 0.05). All the statistical tests were done in R. We used customized perl script for all the analysis used in this study. We used R ggplot2 library for plotting (12).

### Cell culture

U2OS, HDFF, HEK293T cells were cultured in Dulbecco’s Modified Eagle Medium (#10313-021, Gibco) and Merkel Cell Carcinoma cell lines, MKL-1 and MS-1, were cultured in RPMI Medium (#11875-093, Gibco). All culture mediums contained 10% fetal bovine serum, 1X Penicillin/Streptomycin (#15140-122, Gibco) and 1X GlutaMAX -1 (#35050-061, Gibco). Cells were maintained at 37°C and 5% CO2.

### Stem-loop reverse transcription-Real time polymerase chain reaction (RT-qPCR)

#### First strand synthesis

The amplification of piRNA and mature microRNAs was carried out using a stem-loop gene-specific reverse transcription primer in a two-step reaction. Briefly, 100 ng of total isolated RNA and 1 pmol stem-loop primer were added and incubated at 80 °C for 5 min and then at 50 °C for 5 min following which a master mix containing 2.0 µl 10× RT Buffer, 1.0 µl 10 mM dNTP mix, 1.0 µl 0.1 M DTT, 4.0 µl 25 mM MgCl_2_, 0.5 µl RNase OUT (40U/µl) and 0.5 µl SuperScript III Reverse Transcriptase (#55066, Invitrogen, 200 U/µl) was added to make a final volume of 20 μl. The cDNA synthesis was carried out at 25 °C for 5 min, followed by 50 °C for 1 h, and finally, 75 °C for 15 min for enzyme inactivation. This was followed by addition of 0.5 μl RNase H (2U/μl) and incubation at 37°C for 20 min. Stem-loop RT primers (piR62011, 5’ -CTCAACTGGTGTCGTGGAGTCGGCAATTCAGTTGAGACCTCACC-3’; miR182, 5’-CTCAACTGGTGTCGTGGAGTCGGCAATTCAGTTGAGAGTGTGAG-3’; miR183, 5’-CTCAACTGGTGTCGTGGAGTCGGCAATTCAGTTGAGAGTGAATT-3’; miR96, 5’-CTCAACTGGTGTCGTGGAGTCGGCAATTCAGTTGAGAGCAAAAA-3’ and U6, 5’-CTCAACTGGTGTCGTGGAGTCGGCAATTCAGTTGAGAAAATATG-3’) were designed to specifically reverse transcribe mature miRNAs of interest. First-strand cDNA was stored at -20°C until qPCR analysis.

### Real Time quantitative PCR

qPCR was then performed on cDNA using SYBR green (#K0221, Thermo) on a ViiA™ 7 Real-Time PCR System (Applied Biosystem) with gene-specific primers: piR62011 & miR182, F, 5’-ACACTCCAGCTGGGTTTGGCAATGGTAG-3’; miR183, F, 5’-ACACTCCAGCTGGGTATGGCACTGGT-3’; miR96, F, 5’-ACACTCCAGCTGGGTTTGGCACTAGC-3’ U6, F, 5’-ACACTCCAGCTGGGGTGCTCGCTTCG-3’; and Universal Reverse primer, 5’-TGGTGTCGTGGAGTCGCAATTCAGTTG-3’. The following thermal cycling conditions were used: 50°C for 2 min, 95°C for 10 min, 5 cycles of 95°C for 15 s, 45°C for 30 s and 72°C for 15 s followed by 35 cycles of 95°C for 15 s, 55°C for 30 s and 72°C for 15 s and melt curve. PCR products were resolved on a 10% non-denaturing acrylamide gel and stained with ethidium bromide in 1× Tris borate-EDTA.

### TOPO cloning

The amplified cDNA fragments were cloned into the TA cloning vector pCR2.1 (#45-0641, Invitrogen) and the nucleotide sequence was determined. Reaction mixture containing 2 µl PCR product, 1 µl salt solution and 0.5 µl PCR 2.1 TOPO vector to a total volume of 6 µl with nuclease-free water was incubated at RT for 30 min and kept on ice for 2 min followed by transformation in *E*.*coli* XL-10. Plasmid extraction was carried out using plasmid miniprep kit (#27106, Qiagen), following the manufacturer’s protocol, and was sequenced using M13, F, 5’-GTAAAACGACGGCCAG-3’ and M13, R, 5’-CAGGAAACAGCTATG-AC-3’ primers.

### Standard curve generation

To plot a standard curve of Copy numbers to Ct values, the concentrations of piR62011, miR183, miR182, miR96 and U6 cDNA cloned in PCR2.1 TOPO vector was determined using Nanodrop 2000 (Thermo Fisher) and its subsequent copy number was calculated using NEB dsDNA calculator. The plasmids were diluted from 1 copy to 10^8^ copies and amplified using SYBR Green. A standard curve was plot for Ct value obtained corresponding to Copy Number calculated.

### Western blotting

Whole Cell lysates were prepared from the indicated cell lines in EBC lysis buffer (50 mM Tris HCl pH-8, 150 mM NaCl, 0.5mM EDTA, β-mercaptoethanol-1:10,000, 0.5% NP-40, and protease inhibitor (#5892953001, (cOmplete) Roche Applied Science), incubated for 20 min on ice. The lysate was centrifuged at 14000 rpm for 10 min at 4°C. Protein concentrations were quantitated using a standard BCA assay, and samples were resolved on a 12% SDS-PAGE gel. Prior to loading, the protein sample was mixed with 5X Laemmli buffer and incubated at 99°C for 10 min, cooled immediately on ice. The protein was transferred onto a PVDF membrane (#162-0177, Biorad), then blocked in 5% non-fat milk in 0.1% Tris-buffered saline-Tween 20 (TBST) at room temperature for 1 h. All primary antibodies were incubated with the membrane at 4 °C overnight at the following concentrations: Ab3 (1:1000), Ab5 (1:1000), Vinculin (1:15000) (#V9131, Sigma). Membranes were washed thrice for 15 min and incubated with the secondary antibody (sheep anti-mouse HRP, 1:5000) for 1 h at room temperature. After three more washes, the protein bands were detected with the ECL Western blotting Detection System (#RPN2232, GE Healthcare) and recorded on ImageQuant™ Las 4000 (GE Healthcare). Image J software was used to quantify band intensities.

### Luciferase Reporter Assays

U2OS cells were transiently cotransfected with 5 µg of the firefly luciferase reporter plasmid (pGL4-miR182 promoter), 5 μg of early region of MCV genome-expressing plasmid (ER21) or pLenti6.3 empty control vector and pRL.TK (Renilla luciferase) using Lipofectamine 3000 Transfection reagent (# L3000015, Invitrogen) according to the manufacturer’s directions. Cells were harvested 48 h post-transfection using a plastic cell scraper, lysed in 1× Passive Lysis buffer (#E1910, Promega), and incubated at RT for 15 min on a rotor disk. The lysate was assayed for firefly and *Renilla* luciferase activity following the Promega protocol using a luminometer. Firefly luciferase activity was normalized to corresponding Renilla luciferase activity.

### Surface Plasmon Resonance

SPR single-cycle kinetics was performed to study the interaction of CMBLs with pre-miR182. The experiment was carried out in a BIAcore T200 platform by immobilizing 5’-biotinylated pre-miR182 (RU 557) onto a series S sensor chip SA surface. 5’ Biotinylated pre-miR182 (0.2 μM) in HBS-N running buffer [0.01 M, HEPES {4-(2-hydroxyethyl)-1-piperazineethanesulfonic acid buffered saline} and 0.15 M NaCl, pH 7.4] was immobilized onto a SA sensor chip by avidin-biotin coupling in flow cell 2 at a flow rate of 5 μl/min, 60 sec of contact time with an immobilization level of 500 RU. Blank immobilization was performed in the flow cell 1 to permit reference subtraction. Ligand solution (1 mM DMSO stock) was diluted using HBSEP+ buffer [0.01 M HEPES, 0.15 M NaCl, 3.0 mM EDTA, pH 7.4, 0.005% (v/v) Surfactant P20] to a final stock concentration of 10 µM in 1x HBSEP+ buffer containing 5% DMSO. Final running concentrations of ligand were prepared using 10 µM stock such that all final ligand solutions were contained 5% DMSO in 1x HBSEP+ buffer. Sensorgrams were obtained in the concentration range of 0.25–4.0 µM with 60 mL/min flow rate, 30 min of contact time and 120 sec of dissociation time. All sensorgrams were corrected by reference subtraction of blank flow cell response and buffer injection response.

### Lentiviral production and T antigen knockdown

MCV T antigens were knocked down in MKL-1 cells using shRNAs previously published (Houben et al, 2010 and Shuda et al, 2011). The shRNAs were cloned into pLKO.Puro vectors, at Age1 and EcoR1 sites and lentivirus was generated in 293T cells using psPax2 and pVSV.G vectors and MKL-1 cells were infected using spinoculation (centrifugation at 800 ×g for 30 min with viral supernatants) followed by infection overnight in the presence of 1 μg/mL Polybrene. 24 hours post infection, MKL1 cells were spun down and resuspended in medium containing puromycin (1 μg/mL). Cells were harvested after 72 hours and processed for immunoblotting and immunoprecipitation. (sh.panT-AAGAGAGGCTCTCTGCAAGCT, sh.Scr –TAAGGTTAAGTCGCCCTCG).

### Lentivirus based overexpression

Packaging (psPAX2) and envelope (pVSV.G) plasmids were co-transfected with the overexpression construct into HEK 293T cells using Lipofectamine 3000. Two days after transfection, 293T cell supernatant was purified with 0.45 μm filter and concentrated using Amicon Ultra centrifugal filter (#UFC903024, Sigma) followed by centrifugation at 4000×g for 15 min. The lentiviral concentrate was mixed with Lenti-X concentrator (#631231, Clontech) to increase the viral titre, incubated for 2 hrs and spun at 1500 ×g for 45 min at 4°C. The viral pellet was resuspended in fresh media supplemented with 1 μg/ml polybrene while transducing recipient HDFF cells. Stable cell lines were generated after selection with 1 μg/ml puromycin.

### Immunoprecipitation followed by RNA extraction

MKL-1 cell lysate was prepared in RIPA buffer (#R0278, Sigma) 150 mM NaCl, 1.0% IGEPAL, CA-360, 0.5% sodium deoxycholate, 0.1% SDS and 50 mM Tris-HCl and freshly added RNase inhibitor (#M03145, BioLabs), protease inhibitor (#5892953001, (cOmplete) Roche Applied Science) and phosphatase inhibitor (#4906845001, (PhosSTOP) Sigma Aldrich) cocktail, incubated for 20 min on ice. The lysate was centrifuged at 14000 rpm for 10 min at 4°C to remove debris. For comparison, 1/10^th^ of the protein fraction was aliquoted as Input and total RNA was isolated using TRIzol Reagent (Life Technologies). 5 μg of rabbit polyclonal PIWIL4 antibody (#PA5-31448, Thermo) was added to the remaining protein fraction and kept at 4 °C for 2 hrs on a rotating apparatus followed by overnight incubation with 30 μl of the protein G plus agarose beads (#SC-2002, Santa Cruz) at 4 °C. Next day, beads were washed thrice with 1× RIPA buffer containing 0.2U RNase inhibitor and 1X protease inhibitor and the enriched RNA was isolated with TRIzol reagent, in accordance with the manufacturer’s protocol. 250ng of Input and IP were used for RT-qPCR analysis.

## ACKNOWLEDGEMENTS

We thank Professor Thomas Andl from the Vanderbilt University Medical Center, Nashville, USA for sharing sequencing data regarding the miRNome of MCC. We also thank him for discussions and follow up email discussions and support. We also thank Srikar Krishna for excellent scientific discussions that were very helpful. RA would also like to thank Zaina, Ziyah, and Karan for their love, strength, and support.

